# Efficient delivery of gene editors using intein-engineered virus-like particles

**DOI:** 10.64898/2026.01.25.701600

**Authors:** Guannan Zhou, Vicky W.Q. Hou, Houze Zhou, Samantha Roudi, Juliette Suermondt, Oskar Gustafsson, Zheyu Niu, Doste R. Mamand, Lien Van Hoecke, Oscar P. B. Wiklander, Roosmarijn E. Vandenbroucke, Joel Z. Nordin, André Görgens, Samir EL Andaloussi, Xiuming Liang

## Abstract

Virus-like particles (VLPs) represent a promising next-generation drug delivery platform. However, conventional VLPs rely on multiple viral components for effective cargo encapsulation and delivery, raising safety concerns. Here, we present a novel strategy to engineer immature VLPs using a self-cleaving intein system. We employed viral Gag proteins as sorting domains, linking cargo proteins to Gag through inteins, thereby eliminating the need for the conventional protease cleavage typically mediated by the gag-pol protein. During VLP biogenesis, intein-mediated cleavage released cargo proteins into the lumen, enabling efficient intracellular delivery when VLP surfaces are pseudotyped with VSV-G. Optimal candidates for delivering Cre recombinase and gene editing tools (Cas9, Cas12a and base editors) were identified by screening various Gag proteins. Notably, these VLPs achieved robust gene editing in primary cells, including naïve and activated T cells, as well as hematopoietic stem and progenitor cells (HSPCs). A single local intracerebroventricular (ICV) infusion of optimized particles induced up to 60% tdTomato expression in the brain regions of reporter mice, while intravenous injection resulted in significant recombination (up to 70%) of a variety of cell types across organs. Collectively, we developed a simplified, efficient VLP platform for intracellular cargo delivery with broad therapeutic potential for gene editing and treatment of human diseases.

## Introduction

Genome editing tools, including CRISPR/Cas9, CRISPR/Cas12a, base editors and prime editors, have gained tremendous interest due to their ability to permanently modulate gene expression^1, 2, 3, 4^. An ever-growing number of delivery approaches have been explored and harnessed to improve their delivery into cells. These strategies can be broadly divided into two main categories: viral and non-viral carriers. Among viral vectors, adeno-associated viruses (AAVs) are the most widely used, showing strong gene-editing performance in both mice and non-human primates (NHPs)^5^. In the non-viral category, lipid nanoparticles (LNPs) are commonly employed to deliver mRNAs encoding the key components of gene-editing tools^6, 7^. Meanwhile, virus-like particles (VLPs) have emerged as one of the most promising new delivery platforms, exhibiting remarkable delivery efficiency in cultured cells and in mice^8, 9, 10^.

To engineer VLPs, the Gag protein is tethered to cargo proteins and serves as a scaffold for packaging them into the lumen. A cleavable peptide, recognized and cut by a protease, is typically used to link the Gag protein and cargo, allowing the cargo to be released after co-transfection with the gag-pol protein, which produces the necessary protease^8, 9, 11^. However, gag-pol also encodes integrase, reverse transcriptase, and other components, raising safety concerns. This highlights a need for alternative strategies to separate cargo from the Gag scaffold without relying on gag-pol–derived proteases.

Apart from cleavable peptides, other separation systems have been developed to enable cargo protein loading and release in related lipid-bilayer vesicles, i.e. exosomes, such as EXPLORs^12^ and MAPLEX^13^. EXPLORs (Exosomes for Protein Loading via Optically Reversible protein–protein interactions) utilize a blue light-controlled, reversible protein–protein interaction module integrated with the endogenous process of exosome biogenesis. Upon blue light illumination, cargo proteins are recruited and loaded into developing exosomes, and once the light is removed, the interaction is reversed, resulting in the release of cargo proteins into the intraluminal space of the exosomes. In contrast, MAPLEX (mMaple3-mediated protein loading into and release from exosomes) employs another photocleavable protein, mMaple3, fused to exosomal membrane markers and cargo proteins. Upon blue light exposure, photocleavage is induced, enabling the detachment and subsequent release of the cargo proteins from the exosomal membrane into the lumen. However, these separation systems are dependent on light-control, which makes them more complex and increases production-and operational costs.

To simplify the engineering of traditional VLPs, we utilized a self-cleaving protein, intein, previously used by us for engineering of exosomes^14^. The intein was inserted between the Gag protein and the cargo, enabling the release of soluble cargo into the lumen during VLP biogenesis. By incorporating the fusogenic protein VSV-G to enhance endosomal escape in recipient cells, we achieved highly efficient intracellular delivery of different biotherapeutics. In our study, we evaluated six viral Gag proteins (FMLGag^8^, HV190Gag^15^, HV1N5Gag^16^, MMLGag^9, 17^, BaEVGag^18^ and HFVGag^19^) and two commonly used human Gag homologs (hArc^20^ and hPEG10^21^, supplementary Table 1). Among the tested candidates, two Gag proteins, FMLGag and HV190Gag, were identified as optimal for delivering Cre recombinase and gene-editing tools (CRISPR/Cas9^22^, CRISPR/Cas12a^23^ and base editors^24, 25^), respectively. In short, intein-engineered VLPs represent a programmable platform capable of delivering a wide range of biotherapeutics, offering significant potential for the development of next-generation VLP-based therapies with enhanced efficacy.

## Results

### Intein-engineered VLPs enable efficient intracellular delivery of Cre recombinase

To explore the potential of engineering immature VLPs with an intein, we evaluated several published Gag proteins from different viruses to identify the most effective scaffolds for delivery (Fig. 1a and 1b, and Supplementary Table 1). Engineered VLPs were isolated using tangential flow filtration (TFF)^14, 26^ (Fig. 1c). The isolated VLPs were characterized by nanoparticle tracking analysis (NTA), which revealed a size distribution between 90 nm and 120 nm (Supplementary Fig. 1a). Transmission electron microscopy (TEM) confirmed that the engineered VLPs exhibited a typical spherical morphology surrounded by a host cell–derived lipid bilayer envelope (Supplementary Fig. 1b). This bilayer, acquired during budding, incorporates the engineered viral glycoproteins such as VSV-G, consistent with the structural features of enveloped retrovirus-like particles^27, 28^.

**Fig. 1.**
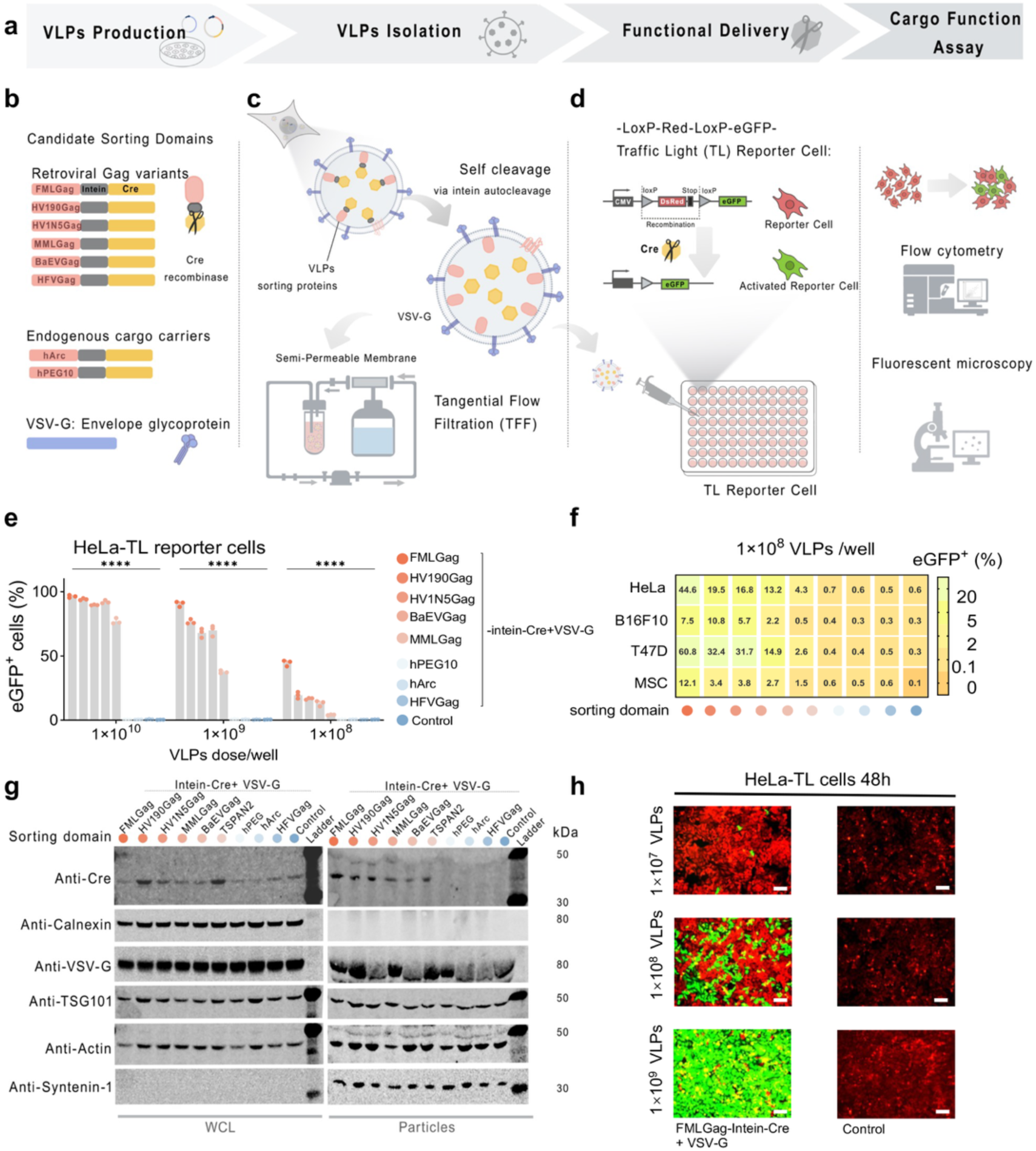
Identification of efficient intein-engineered VLPs scaffold proteins for functional intraluminal cargo delivery. **(a)** Overview of the experimental workflow, including VLPs production, isolation, delivery, and cargo function assay using reporter cells. **(b)** Design of candidate sorting domains fused to Cre recombinase via an intein-based self-cleavable linker. Categories include retroviral Gag variants, human Gag homologs (e.g.hArc, hPEG10) and control VSV-G. **(c)** Schematic of VLPs production by transient transfection and purification via tangential flow filtration (TFF). Intein-mediated autocleavage allows luminal release of Cre cargo. **(d)** Functional delivery assay using “Traffic Light” (TL) reporter cells containing a LoxP-Red-LoxP-eGFP cassette. Cre delivery induces recombination and eGFP expression, which is detectable by flow cytometry or fluorescence microscopy. **(e)** Quantification of eGFP⁺ HeLa-TL cells after 48 h incubation with VLPs containing different sorting domains at serial doses (1×10^10^, 10^9^, 10^8^ particles/well). All dose–response experiments involving engineered VLPs in cultured cells were carried out in a 96-well plate format, unless indicated otherwise. **(f)** Functional delivery efficiency (eGFP⁺ %) of selected VLPs (10^8^ particles/well) in four TL reporter cell types: HeLa-TL, B16F10-TL, T47D-TL, and MSC-TL. **(g)** Western blot analysis was performed using lysates from 5×10^5^ particle-producing cells and 1×10^10^ engineered particles respectively. TSG101, syntenin-1, and β-actin served as particle markers, while calnexin (an endoplasmic reticulum marker) was included to confirm absence of cellular contaminants in particle preparations. Control is the Intein-Cre+VSV-G group. **(h)** Representative immunofluorescence images of HeLa-TL cells treated with FMLGag-Intein-Cre +VSV-G VLPs or control particles at indicated doses. GFP activation indicates successful delivery. Control is the Cre-loaded particles without VSV-G pseudo tying group. Scale bar, 100 μm. *Statistical significance: ****p < 0.0001*.

Cre recombinase was selected as the model cargo, and functional activity was assessed using traffic-light (TL) reporter cells (Fig. 1d), as described previously^14, 29^. In this assay, recombination mediated by Cre activates eGFP expression, which can be quantified by flow cytometry or visualized via fluorescence microscopy.

Across multiple reporter cell lines (HeLa-TL, T47D-TL, B16F10-TL, and MSC-TL), the engineered VLPs induced dose-dependent eGFP activation (Fig. 1e and 1h, Supplementary Fig. 2a-2c, Supplementary Fig. 8a). Five Gag candidates, FMLGag, HV190Gag, HV1N5Gag, BaEVGag, and MMLGag, achieved substantial eGFP expression with as few as 1×10^8^ particles/well (Fig. 1f and Supplementary Fig. 2a-2c). Notably, the best-performing candidate, FMLGag, mediated notable Cre-dependent recombination in HeLa-TL and MSC-TL cells even at 1×10^7^ particles per well (Supplementary Fig. 2d-2g). However, not all Gag proteins were effective scaffolds for cargo loading. Gag from human foamy virus (HFVGag) failed to mediate detectable Cre transfer in any reporter cell line (Fig. 1e and 1f), underscoring the importance of screening for functional scaffolds. Likewise, two human Gag homologues, hPEG10 and hArc (Supplementary Table 1), showed no measurable Cre delivery (Fig. 1e and 1f). To investigate the basis of these differences, we performed Western blot (WB) analyses on whole cell lysates (WCL) and isolated particles. Cre was expressed and cleaved in all producer cells; however, cleaved Cre was detected in isolated particles only for the six functional scaffolds (FMLGag, HV190Gag, HV1N5Gag, BaEVGag, MMLGag, and TSPAN2, which served as a positive control^28^) (Fig. 1g). These results indicate that the efficiency of Cre delivery depends on the inherent cargo-loading capacity of the scaffolds. In addition, we directly compared the delivery efficiency of Cre by intein-engineered VLPs with the published Nanoblade system^8^ and found significant improvement over the latter (Supplementary Fig. 3a and 3b). Because the Gag protein lacks transmembrane domains and behaves like a cytoplasmic protein, we questioned whether releasing the cargo via intein-mediated cleavage, or other cleavable linkers, would offer a functional advantage. For this, we compared three designs: direct fusion of Cre to FMLGag (FMLGag–Cre); Intein-mediated cleavage using the wild-type intein (FMLGag–Intein–Cre); Cleavage-deficient intein (N440A variant) as C-terminal cleavage–impaired (Supplementary Fig. 4a and 4b). In B16F10-TL and MSC-TL cells, the intein-cleavage design (FMLGag–Intein–Cre) showed a clear advantage over the direct-fusion design, while in HeLa-TL and T47D-TL cells the efficiency was comparable between the two (Supplementary Fig. 4c and 4d). In contrast, the cleavage-deficient intein variant dramatically reduced Cre transfer in all reporter cell lines (Fig. Supplementary Fig. 4c and 4d). These findings indicate that intein-mediated cargo release is generally more effective than non-cleavage designs, possibly because it fully releases the cargo from the scaffold and avoids potential steric or conformational interference caused by direct fusion.

**Fig. 2.**
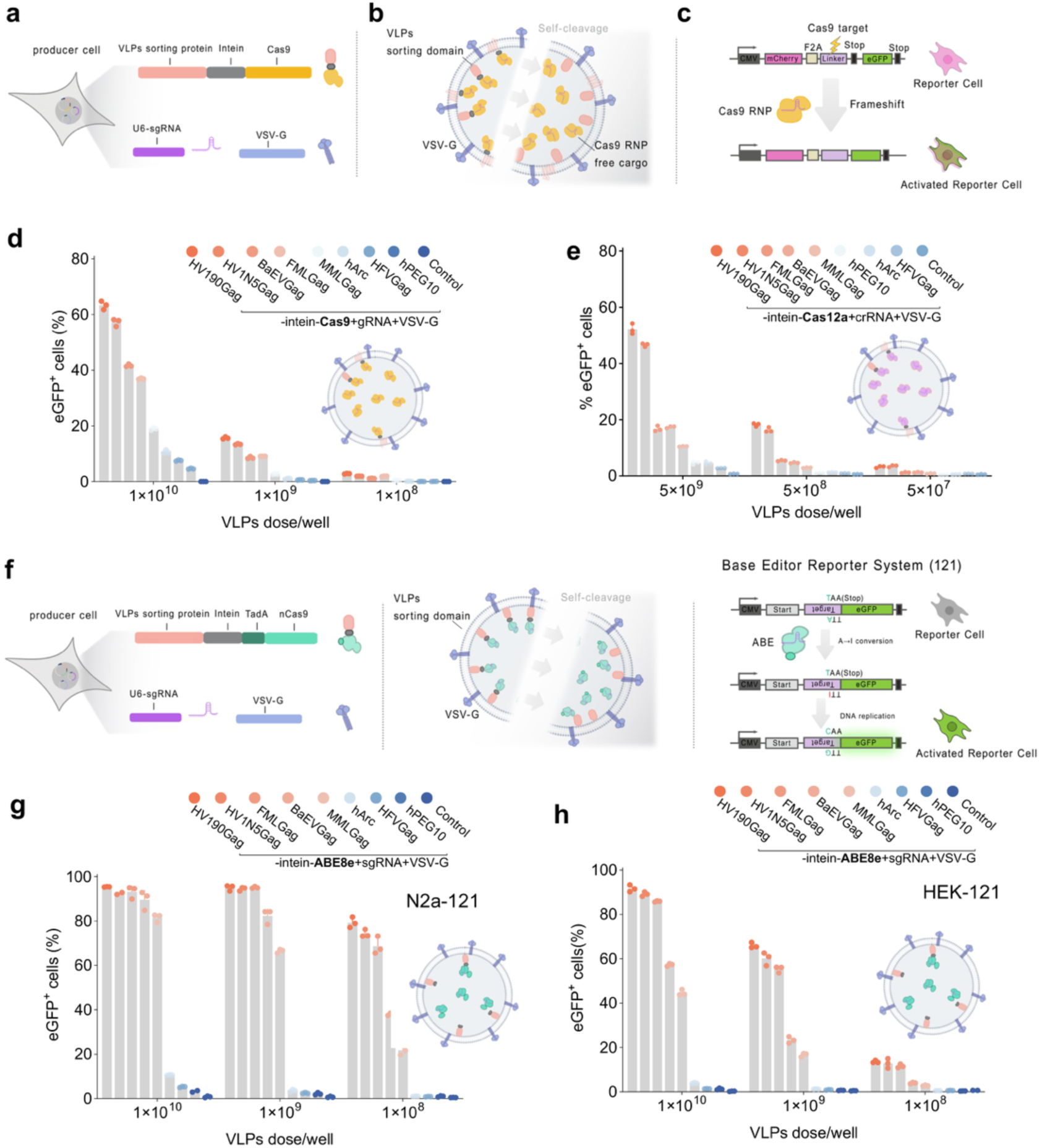
**Fig. 3 Evaluation of intein-engineered VLPs for functional delivery of gene editors. (a)** Schematic of the production system for Cas9 RNP–loaded VLPs. Producer cells were co-transfected with plasmids encoding VLP sorting fusion proteins (Sorting domain–Intein–Cas9), a U6-driven sgRNA construct, and VSV-G. **(b)** Illustration of VLP structure and self-cleavage mechanism. Upon intein-mediated release, Cas9 RNP is packaged inside VLPs for delivery. **(c)** Schematic of the “Stop Light (SL)” reporter cell line, where Cas9-mediated indel at the target site restores eGFP expression through frameshift repair. **(d)** Quantification of eGFP⁺ cells in HEK-SL reporter cells 72 h after treatment with Cas9 RNP VLPs produced using different sorting domains. Efficiency was dose-dependent. Control group is the Cas9 RNP-loaded particles without VSV-G pseudo typing. **(e)** Quantification of eGFP⁺ in Cas12a reporter cells 72 h after exposure to Cas12a RNP VLPs generated with different sorting domains and the editing efficiency was dose-dependent. Control group is the Cas12a RNP-loaded particles without VSV-G pseudo typing. (**f**) Schematic production system for ABE–loaded VLPs and representation of the A→G base-editing reporter. A→G base editing of the antisense strand followed by DNA replication converts the stop codon, TAA, to CAA (glutamine) releases the translation of its downstream eGFP. **(g, h)** Quantification of eGFP⁺ cells in N2a-121 and HEK-121 reporter cells respectively 72 h after treatment with Base Editor (ABE8e) RNP VLPs produced using different sorting domains with serial doses. Control group is the base editor-loaded particles without VSV-G pseudo typing. Data are presented as mean ± SD.

**Fig. 3.**
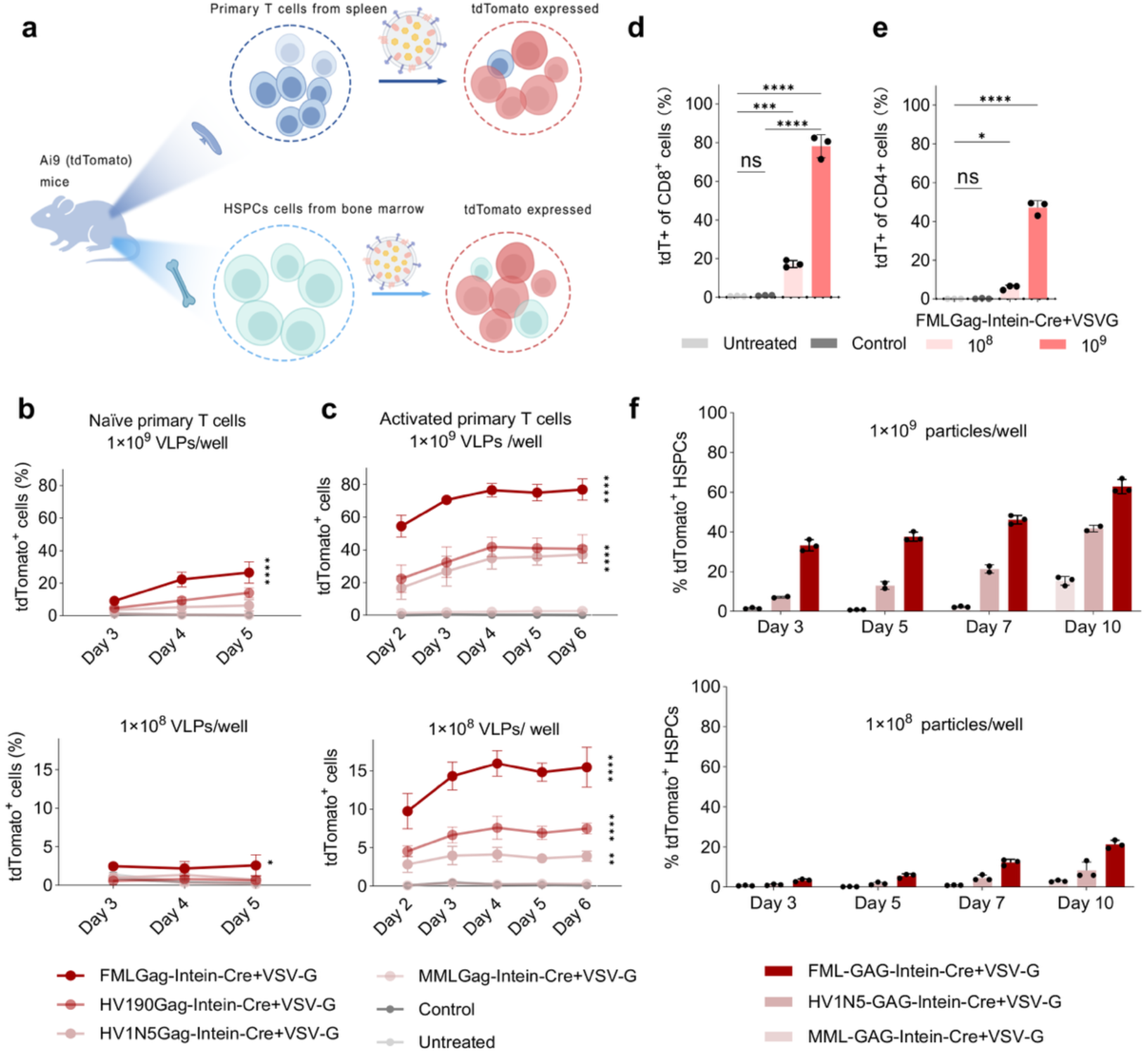
Efficient ex vivo delivery of Cre recombinase into primary cells from Cre-LoxP tdTomato reporter mice. **(a)** Schematic workflow of the ex vivo assays: VLPs were incubated with primary T cells and HSPCs derived from R26-LSL-tdTomato mice. Successful Cre delivery induces tdTomato expression. **(b)** Time-course analysis of naïve tdTomato⁺ T cells treated with VLPs at two doses (1×10^9^ and 1×10^8^ particles/well), measured at 72, 96, and 120 h post-incubation. Control group is the Cre-loaded particles without VSV-G pseudo typing. (c) Time-course analysis of activated tdTomato⁺ T cells treated with VLPs at indicated two doses. **(d, e)** Flow cytometry quantification of activated tdTomato⁺ CD8⁺ (d) and CD4⁺ (e) T cells after treatment with indicated VLPs at day 6. **(f)** tdTomato expression in isolated HSPCs from R26-LSL-tdTomato mice after treatment of engineered particles at different time points. Data are presented as mean ± SD. *Statistical significance: *p < 0.05, **p < 0.01, ***p < 0.001, ****p < 0.0001; ns, not significant*.

**Fig. 4.**
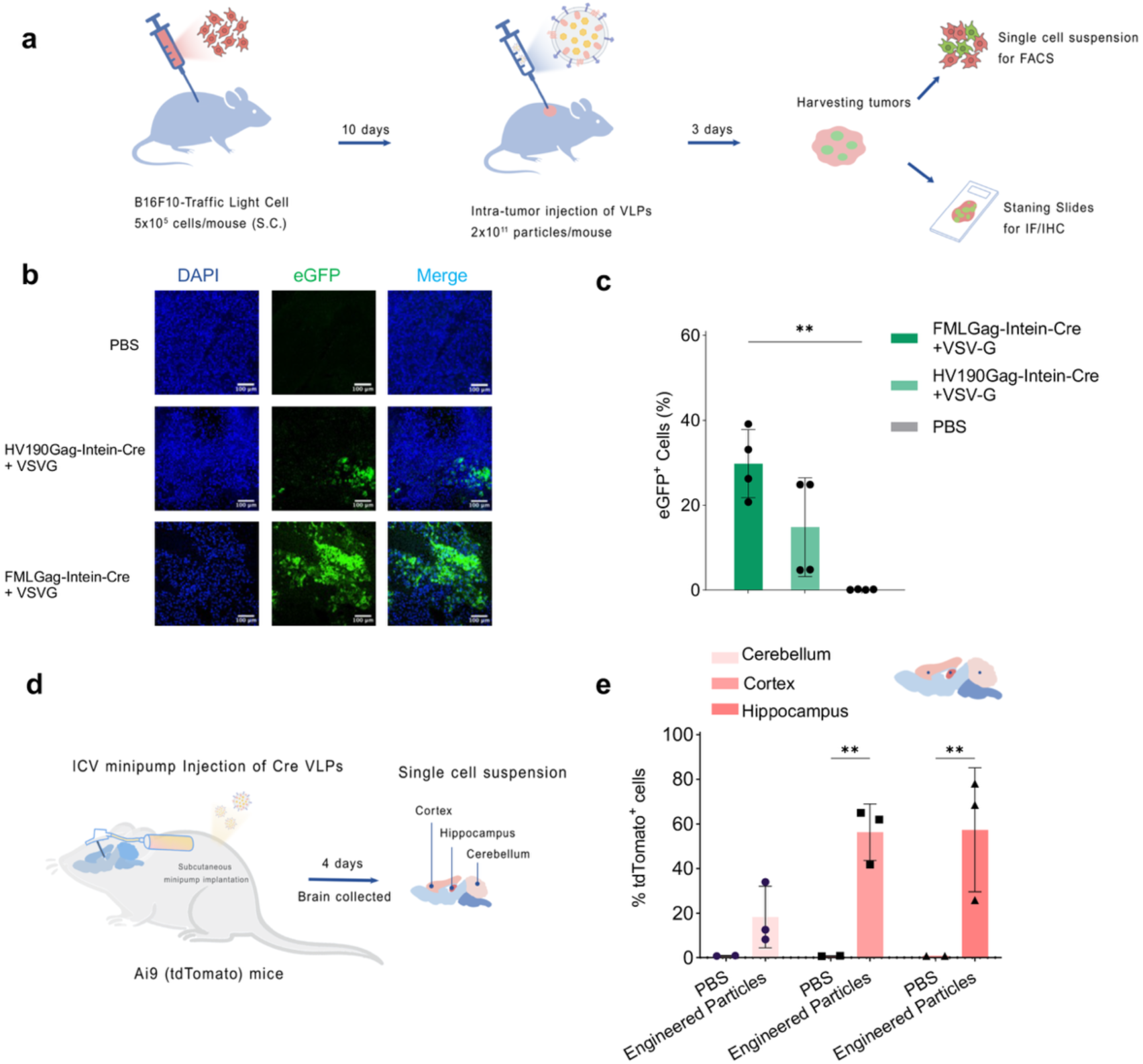
In vivo Cre recombination in tumor and brain tissues by local injection of intein-engineered VLPs. **(a)** Schematic of intra-tumoral VLP delivery: B16F10-Traffic Light (TL) cells were inoculated subcutaneously (5×10^5^ cells/mouse), followed by intra-tumoral injection of Cre VLPs (2×10^11^ particles/mouse) 10 days later. Tumors were harvested 3 days post-injection for flow cytometry and immunofluorescence analysis. **(b)** Representative fluorescence images showing eGFP expression in tumor sections, resulting from CRE-mediated recombination. Scale bar, 100 μm. **(c)** Quantification of eGFP⁺ cells in tumors by flow cytometry. n = 4 mice per group. (d) Schematic of intracerebroventricular (ICV) minipump infusion model: R26-LSL-tdTomato mice were infused with Cre VLPs (8×10^11^ particles/mouse) into the lateral ventricles. Brains were collected 4 days after particle infusion for single-cell dissociation. **(e)** Quantification of tdTomato⁺ cells in different brain regions (cortex, hippocampus, cerebellum) following ICV injection. Cre-loaded VLPs induced detectable recombination across all tested regions. n = 3 mice for engineered particle group and n = 2 mice for PBS group. Data are presented as mean ± SD. *\*\*p < 0.01*.

In summary, we successfully applied intein engineering to VLPs and identified specific Gag proteins with superior capacity to package and deliver functional protein cargoes.

### Intein-engineered VLPs efficiently deliver different gene editors

Following the successful delivery of Cre recombinase, we next evaluated the capacity of intein-engineered VLPs to deliver more therapeutically relevant payloads, specifically, genome-editing tools. Given the broad applicability of CRISPR/Cas9, we replaced the Cre cargo with Cas9 in the intein-engineered VLPs (Fig. 2a). Upon co-transfection with a single guide RNA (sgRNA), the goal was to encapsulate Cas9 ribonucleoproteins (RNPs) into the particles (Fig. 2b) and assess their editing activity using stoplight (SL) reporter cells^30^ (Fig. 2c). In this reporter system, as described previously^14, 30^, Cas9-mediated editing of a linker sequence between mCherry and eGFP activates eGFP expression.

Among the Gag scaffolds tested, HV190Gag exhibited the highest efficiency in inducing eGFP-positive cells, followed by HV1N5Gag and BaEVGag (Fig. 2d, Supplementary Fig. 5, Supplementary Fig. 8b). Interestingly, FMLGag, which was the most effective for Cre delivery, showed only modest activity in delivering Cas9 RNPs (Fig. 2d), underscoring the importance of screening Gag scaffolds for each specific cargo type. We also compared intein-mediated cleavage with the cleavable peptide approach used in the previously reported Nanoblade system^8^. Western blot analysis of both producer cell lysates and isolated particles revealed that intein achieved more efficient cleavage of Cas9 (Supplementary Fig. 6a).

To extend these findings, we evaluated delivery of another widely used genome-editing tool, CRISPR/Cas12a^31^, using the same intein-engineered VLP platform. The Cre or Cas9 cargo was replaced with Cas12a, and editing was measured in SL reporter cells containing a linker sequence specifically targeted by the Cas12a/crRNA complex. The results again identified HV190Gag and HV1N5Gag as the most effective scaffolds (Fig. 2e), consistent with the Cas9 data (Fig. 2d). We next extended the platform to deliver base editors and verified precise base conversion (Fig. 2f). Using previously described reporter cells, an A→G transition removes a stop codon upstream of eGFP, enabling fluorescence detection by flow cytometry^32^. Intein-engineered VLPs mediated highly efficient delivery of adenine base editor ABE8e, with HV190Gag performing best, followed by HV1N5Gag, FMLGag, BaEVGag, and MMLGag (Fig. 3g-3h, Supplementary Fig. 6b). In N2a-121 cells, as few as 1×10^8^ VLPs yielded close to 80% eGFP activation (Fig. 2g). Comparable dose-dependent editing was observed in HEK-121 and T47D121 cells (Fig. 2h, Supplementary Fig. 6b).

In summary, intein-engineered VLPs can efficiently package and deliver multiple classes of genome-editing tools, including Cas9 RNPs, Cas12a RNPs and base editors.

### Efficient ex vivo gene editing of mouse primary T cells and HSPCs

To further evaluate whether Gag-engineered VLPs can functionally deliver cargo into transfection refractory primary T cells^35, 36^, we isolated T cells from R26-LSL-tdTomato reporter mice (Fig. 3a). These mice carry a LoxP–STOP signals–LoxP cassette between the CAG promoter and the open reading frame (ORF) of the red fluorescent protein tdTomato in the ROSA26 locus^14, 33, 34^. In the presence of Cre recombinase delivered by VLPs, the STOP cassette is excised, enabling tdTomato expression in recipient cells. Spleens were harvested from mice, and CD3⁺ primary T cells were isolated for treatment with Cre-loaded VLPs (Fig. 3a). Cells were divided into two groups: activated T cells (primed with CD3/CD28) and naïve T cells (no priming). FMLGag-engineered VLPs edited both activated and naïve T cells in a dose-and time-dependent manner (Fig. 3b,c and Supplementary Fig. 8d). Across incubation times ranging from day 2 to day 6, editing efficiency peaked at around 80% in activated T cells when treated with 1×10^9^ particles per well for 96 h (Fig. 3c). In naïve T cells, editing reached about 30% with 1×10^9^ particles at day 5 (Fig. 3b). To evaluate editing efficiency in specific T cell subsets, activated T cells treated with FMLGag-VLPs were stained for CD4 and CD8 markers at day 6. Cre recombinase delivery was highly efficient, achieving up to 80% editing in the CD8⁺ population and nearly 50% in CD4⁺ cells (Fig. 3d and 3e).

In addition to primary T cells, we isolated HSPCs from R26-LSL-tdTomato reporter mice^35^ and assessed Cre transfer using intein-engineered VLPs (Fig. 3a). VLPs built with FMLGag, HV1N5Gag, or MMLGag produced robust, time-and dose-dependent Cre-mediated recombination, with FMLGag performing best (Fig. 3f).

### Cre-mediated recombination in melanoma-xenografts and in the central nervous system (CNS) following local VLP administration in mice

Based on in vitro and ex vivo results, we next evaluated the in vivo applicability of the platform using two local administration routes in mice. The first involved intratumoral (IT) injection into B16F10-TL tumor-bearing mice (Fig. 4a), and the second involved intracerebroventricular (ICV) infusion in R26-LSL-tdTomato reporter mice using osmotic minipumps (Fig. 4d).

In the intratumoral model, C57BL/6 mice bearing subcutaneous B16F10-TL melanoma xenografts received direct IT injections of engineered VLPs and three days post-injection, tumors were harvested for paraffin section preparation and single-cell suspensions, respectively. Immunofluorescence (IF) revealed strong eGFP signals in tumors treated with FMLGag-engineered VLPs (Fig. 4b), confirming efficient delivery of bioactive Cre recombinase in vivo. Flow cytometry analysis showed that VLPs treatment achieved over 30% editing of the evaluated tumor cells (Fig. 4c, Supplementary Fig. 8c). In the ICV-minipump model, R26-LSL-tdTomato reporter mice received a continuous infusion of FMLGag-engineered particles for 24 h into the ICV space. 4 days after infusion, single-cell suspensions from different brain regions were analyzed by flow cytometry. Remarkably, we observed nearly 60% recombined tdTomato-positive cells in the cortex and hippocampus, and up to 20% in the cerebellum (Fig. 4e and Supplementary Fig. 7). To our knowledge, this represents the highest reported levels of in vivo brain editing using non-viral delivery, highlighting the strong potential of intein-engineered particles as a delivery platform for therapeutic applications in the CNS.

### Efficient systemic delivery using intein-engineered VLPs

Following local IT and ICV studies, we next evaluated a systemic administration route, intravenous (i.v.) injection, using the same reporter mice with a single-dose regimen (Fig. 5a). The FMLGag-engineered VLPs enabled broad Cre delivery across multiple organs. In bone marrow, editing was detected in approximately 1.5% of total CD45⁺ leukocytes. Within the edited myeloid compartment, nearly half of CD11b⁻ macrophages were tdTomato-positive, whereas other myeloid subsets remained largely unedited. Among lymphoid populations in bone marrow, editing was most pronounced in NK cells (∼40%), with lower but detectable levels in plasmacytoid dendritic cells (pDCs) and T cells (∼5%), showing a modest enrichment in CD4⁺ T cells (Fig. 5b; Supplementary Fig. 9).

**Fig. 5.**
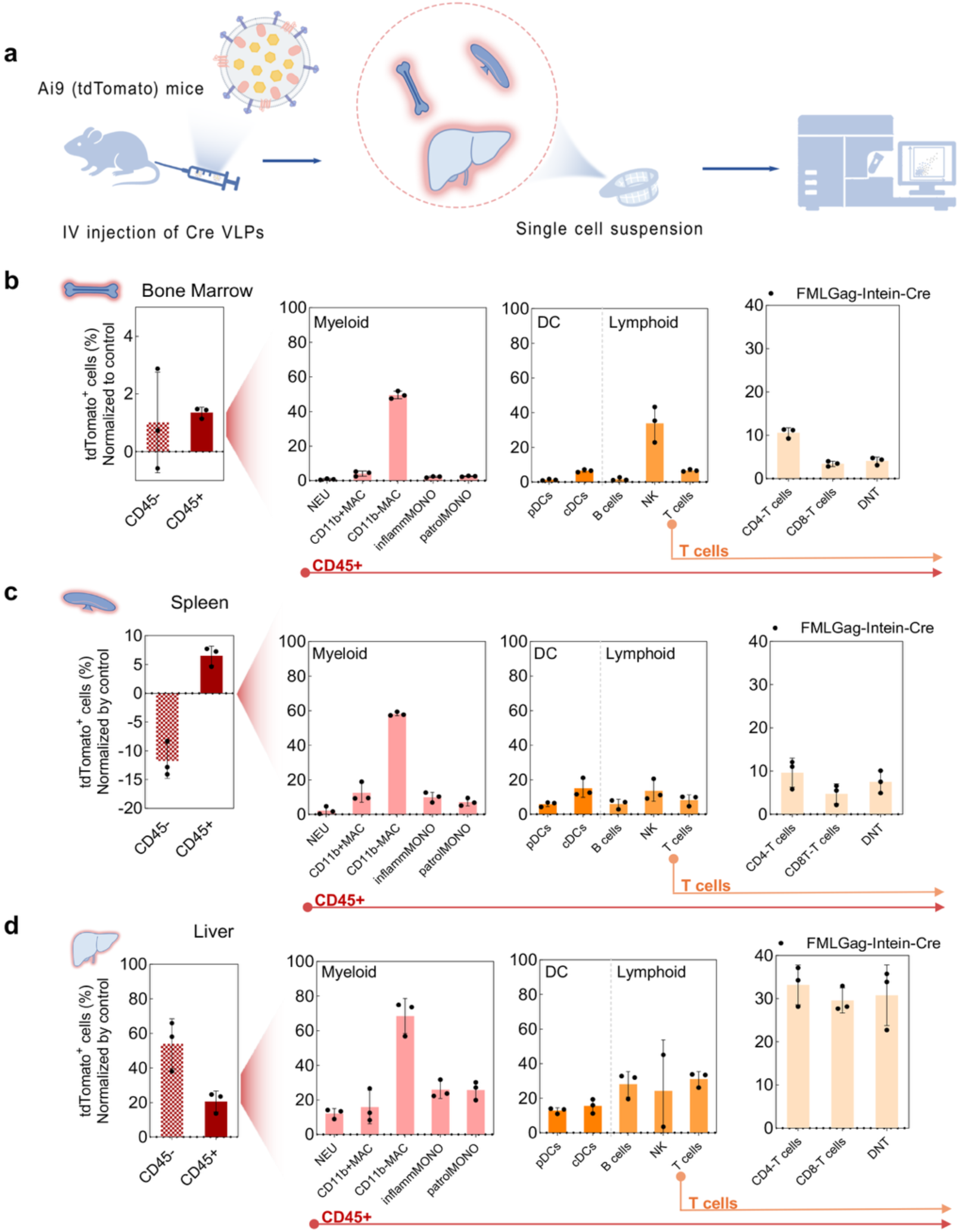
In vivo Cre recombination in vivo by systemic injection of intein-engineered VLPs. **(a)** Schematic of intein-engineered VLPs systemic delivery: intra-venous injection of intein-engineered Cre VLPs (1×10^12^ particles/mouse). Bone marrow, spleen and liver were collected 4 days later for single-cell dissociation. **(b-d)** Quantification of tdTomato⁺ cells in diverse subpopulations in bone marrow (b), spleen (c) and liver (d) by spectral flow cytometry. The percentage of tdTomato⁺ cells was normalized by background subtraction of background signals from the PBS control group. n = 3 mice per group. *See supplementary* figure 9 *for gating details*.

In the spleen, editing reached approximately 7% of CD45⁺ leukocytes. Within the myeloid lineage, CD11b⁻ macrophages exhibited high editing efficiencies approaching 60%. Editing was also observed across multiple lymphoid subsets, ranging from 5% to 15% (Fig. 5c; Supplementary Fig. 9).

The liver displayed the most robust editing activity (Fig. 5d). Nearly 60% of hepatocytes (CD45⁻ liver cells) were tdTomato-positive, along with approximately 20% of total CD45⁺ hematopoietic cells. Within the myeloid compartment, editing reached ∼70% in CD11b⁻ macrophages and exceeded 10% in neutrophils, CD11b⁺ macrophages, and inflammatory monocytes, with patrolling monocytes reaching up to ∼30%. Among liver-resident lymphoid populations, B cells and NK cells exhibited ∼30% and ∼25% editing, respectively, while pDCs and classical DCs showed ∼20% editing. Total T cells reached nearly 35% editing, with CD4⁺, CD8⁺, and double-negative T-cell subsets each approaching ∼30%, indicating efficient editing across liver-resident lymphocyte populations (Fig. 5d; Supplementary Fig. 9).

In addition, we evaluated the potential toxicity of the injected particles by analysing the histology of liver. Compared to the PBS group, the particle-treated mice showed no overt tissue injury or inflammatory changes, indicating that systemic administration of engineered VLPs did not elicit severe toxicity (Supplementary Fig. 10).

Together, these results show that engineered VLPs can efficiently deliver protein cargo in vivo following systemic administration, supporting their potential for future clinical translation.

## Discussion

Realizing the full potential of gene therapy will require a diverse toolkit of delivery modalities capable of efficiently encapsulating and transporting sufficient payloads to target cells. Engineered VLPs^37, 38^, which package proteins of interest while lacking viral genetic material, have emerged as a promising platform for delivering gene-editing tools such as ribonucleoproteins (RNPs). These engineered RNP-VLPs are gaining increasing attention for combining the high delivery efficiency of viral systems with the advantage of avoiding risks linked to viral genome integration and prolonged expression of editing agents. Despite this potential, research on effective VLP-sorting proteins (especially the Gag proteins) and innovative VLP engineering strategies remains relatively limited. In this study, we demonstrate that intein-engineered VLPs can be modularly reprogrammed using a versatile set of components to deliver different payloads, including Cre recombinase, Cas9 RNPs, Cas12a RNPs and base editors.

The use of an intein to separate cargo from a fused tripartite construct streamlines VLP engineering and completely eliminates additional viral components such as protease, integrase, reverse transcriptase, and other enzymes. This design yields immature VLPs that lack replication and integration machinery, offering a clear safety advantage over mature VLPs, which more closely resemble infectious viruses. Structural analyses by cryo-EM and scanning transmission EM have shown that mature virus particle contains approximately 1,200–2,500 Gag proteins, whereas immature particle contains around 5,000 Gag proteins^39^. A recent study found that v5 eVLPs, which are predominantly immature, delivered base editors more effectively than particles with mature capsids^40^. Another study indicated that the capsid is not required for packaging and delivering Cas9 RNP complexes to target cell nuclei^11^. Together, these findings support the use of immature VLPs as promising carriers for gene-editing tools.

In our study, the intein-engineering strategy excludes the gag-pol protein entirely, ensuring that all particles produced are immature and lack proteolytic processing. However, because the intein used here is active at 37 °C, substantial pre-cleavage of cargo can occur before encapsulation during cell culture, reducing the loading capacity of the VLPs, which implies the potential space for the future improvement of the current system.

A key takeaway from our study is that intein-engineered VLPs robustly package and deliver Cas9 and Cas12a RNPs as well as base editors, providing a transient, non-integrating modality. This makes them attractive for ex vivo manufacture of edited T cells and HSPCs, in vivo gene correction or regulation in defined tissues, and high-throughput functional genomics, where a short editing window reduces prolonged nuclease exposure and genotoxic risk. The ability to move seamlessly between nuclease RNPs and base editors also broadens indications, from knockouts to precise edits, without changing the delivery paradigm. Importantly, our head-to-head comparison reveals scaffold-specific performance: different Gag backbones may vary in cargo loading, stability, cell entry, and intracellular release, arguing against a one-size-fits-all solution. This underscores the value of systematic Gag screening (and potentially hybrid other non-viral scaffolds) to match each editor and target cell type. Beyond screening, rational optimization, tuning tropism, endosomal escape, and release kinetics, should further elevate efficacy while maintaining safety. Together, these data position intein-engineered VLPs as a flexible delivery chassis for genome and potential epigenome editors, with clear translational paths provided that manufacturing, biodistribution, and immunogenicity are addressed under standardized quality control.

Although the immature VLPs developed in our study demonstrated excellent delivery efficiency for functional cargoes, they still incorporate two viral components: the Gag protein and the VSV-G envelope glycoprotein. Even though we did not observe significant tissue damage by intein-engineered VLPs in the liver after systemic injection (Supplementary Fig. 10), the presence of these viral elements still raises safety considerations, particularly for potential clinical translation. Gag is a major structural protein of retroviruses^41^, and while it is non-enzymatic and incapable of initiating replication on its own, its close resemblance to native viral architecture may influence immune recognition. VSV-G, meanwhile, is a broad-tropism fusogenic glycoprotein that enables entry into a wide range of mammalian cells^42, 43, 44^. This promiscuous entry capability can be advantageous for research purposes but poses a theoretical risk in clinical settings as this could allow particle uptake by non-target tissues or cell types, thus the possibility of off-target effects would be increased. To address these concerns, further engineering strategies focus on replacing VSV-G with more cell-type–specific fusogens or fusogenic proteins from human endogenous retroviruses (HERVs)^45, 46^, modifying Gag to reduce sequence homology with wild-type viruses or replacing viral Gag with functional human Gag homologs (beyond the hArc^20, 47^ and hPEG10^21^ investigated in this study). Such refinements would strengthen the already favourable safety profile of immature, enzyme-free VLPs and align their design more closely with the stringent requirements for clinical-grade delivery platforms.

Notably, in this study, a single intravenous administration of intein-engineered VLPs produced exceptionally high Cre-mediated recombination across mouse organs, with the liver showing the highest recombination. Comparable efficiencies were observed ex vivo in primary cells, with particularly robust editing in both naïve and activated T cells, as well as in HSPCs. These findings highlight the strong potential of this platform for in vivo T cell–directed genome engineering. In principle, by incorporating a cell-specific targeting moiety onto the surface of the intein-engineered VLPs, such as an antibody fragment, ligand, or engineered receptor-binding domain, it should be possible to confer precise tropism toward defined T cell subsets. This would enable targeted gene editing in vivo with minimal off-target cellular uptake, an approach highly relevant for immunotherapy applications. Furthermore, this strategy is not limited to T cells. By swapping the targeting moiety to match the receptor profile of other immune cells, such as B cells, NK cells, dendritic cells, or HSPCs, the VLPs could be reprogrammed for cell-type–specific delivery in a wide range of therapeutic contexts. Such versatility would dramatically expand the applicability of intein-engineered VLPs, enabling targeted gene correction, functional enhancement, or regulatory element insertion across diverse immune and progenitor cell populations. Importantly, the genome-free and protease/integrase/reverse transcriptase–free nature of these particles provides a favourable safety baseline for in vivo applications, although careful evaluation of biodistribution, immune responses, and long-term safety will be essential before clinical translation.

In summary, the high efficiency observed in both ex vivo and in vivo editing not only validates the performance of the intein-engineered VLP system but also opens a clear path toward customizable, cell-type–targeted genome engineering platforms with broad therapeutic potentials.

## Methods

### Ethical statement

All mouse studies complied with institutional and national regulations and were conducted under approved protocols from the Swedish Board of Agriculture (Jordbruksverket; VLP biodistribution model: 13849-2020; tumor model: 2173-2021) and, where applicable, from the Inflammation Research Center, Ghent University (permit LA1400091/LA2400526).

### Cloning

Codon-optimized DNA sequences coding for the scaffold protein including FMLGag, HV190Gag, HV1N5Gag, MMLGag, BaEVGag, HFVGag, hArc, hPEG10, Tetraspanin2, Cas12a were ordered from IDT (Integrated DNA Technologies, USA). The constructs used in this study were then generated from the ordered fragments through restriction enzyme digestion and subsequent self-ligation. The diverse versions of Gag-Intein-Cre were generated from the VSV-G-Intein-Cre (published by our group) using EcoRI and Kpn2I. Similarly, to synthesize Gag-Intein-Cas9 and Gag-Intein-Cas12a constructs, Gag-Intein-Cre was digested for the backbone to clone Cas9 and Cas12a into respectively. All expression cassettes were confirmed by sequencing. Scaffold protein identifiers are listed in Supplementary Table 2. Plasmids are available from the corresponding author upon request.

### Cell culture

To produce functional EVs, HEK-293T (ATCC, CRL-3216) cells and N2a (ATCC, CCL-131) cells were cultured in DMEM medium (high glucose) supplemented with 10% fetal bovine serum (FBS, Gibco, USA) and 1% anti-anti (Gibco, USA). The HEK-SL cells, HEK-SL2 cells as well as the reporter cell lines (HeLa-TL, B16F10-TL and T47D-TL), were cultured in the same medium and conditions as the HEK-293T cells. MSC-TL were cultured in RPMI-1640 medium supplemented with 10% fetal bovine serum (FBS, Gibco, USA) and 1% Anti-anti (Gibco, USA). Cells were cultured at 37°C in a humidified air atmosphere containing 5% CO_2_.

### Establishing stable reporter cell lines

Lentivirus encoding transgenes of interest were produced in HEK-293T cells according to our previous reports^14^. The corresponding virus particles were added into the cells in six-well plates until approximately 70% confluency was reached and then transduced with lentiviral particles overnight. The transduced cells were then moved to T25 flasks and subjected to puromycin (4 g/ml) for expansion and selection.

### VLPs isolation

The VLPs in this study were produced followed a protocol previously reported by our group^14, 48, 49^. Briefly, the transient transfections of the corresponding plasmids were conducted by using polyethylenimine (PEI, Polysciences) with a ratio of 2:1 (PEI: plasmid). At day 1, HEK-293T cells were seeded into the 15-cm dishes (5 million cells/dish) in full DMEM medium. At day 3, the cells were transfected by 30 *µ*g plasmid (20 *µ*g plasmid of each was used for 2 plasmids co-transfection, while 15 *μ*g of each was used for 3 plasmids.) and 6 hours later the medium was changed to Opti-MEM medium (Gibco, USA) with 1% anti-anti. At day 5 (after 48 h of transfection), the conditional medium was collected and centrifuged of two rounds (700 × g for 5 min and then 2000 × g for 10min) to pre-clear pellet cells and cell debris, followed by the filtration through a 0.22 *µ*m filter system.

The filtered conditioned media were diafiltrated and concentrated to approximately 50 mL by a 300 kDa Tangential flow filtration (TFF, MicroKross, 20 cm^2^, Spectrum laboratories)^14, 26^. The particles were further concentrated until around 500 *µ*L by using an Amicon Ultra-15 spin filter with 100 kDa molecular weight cut-off (Millipore). Finally, the concentrated VLPs were collected in 1.5 mL microcentrifuge MAXYmum recovery tubes (Axygene, USA) and the concentrations were determined using Nanoparticle Tracking Analysis (NTA).

### Transmission electron microscopy (TEM)

A 3 μL aliquot of the sample was deposited onto 400 mesh copper grids (Ted Pella) coated with carbon and reinforced with formvar, which had been glow-discharged using an easiGlow™ system (Ted Pella). The sample was allowed to adsorb for roughly 30 seconds before the excess liquid was gently removed by blotting. Grids were then rinsed with Milli-Q water and subjected to negative staining using 1% ammonium molybdate. Transmission electron microscopy was carried out on a Hitachi HT7700 instrument (Hitachi High-Technologies), operated at 80 kV and fitted with a 4-megapixel Veleta CCD camera (Olympus Soft Imaging Solutions GmbH).

### Nanoparticle Tracking Analysis (NTA)

The VLP samples were diluted with 0.22 µm-filtered PBS before being examined with a ZetaView (Particle Metrix, Germany) device to determine particle sizes and concentrations. Samples were resuspended in PBS to a final volume of 1 mL prior to analysis. Optimal particle concentrations were determined through preliminary assessments aimed at achieving a particle count of 140–200 per frame. Measurement parameters were set using the manufacturer’s default settings optimized for virus-like particles (VLPs). Each sample was analyzed in three independent runs, during which 11 positions per well were scanned, and 30 frames were acquired at each position.

### Western blot

Whole-cell lysates (WCL) were prepared using RIPA buffer containing a protease inhibitor cocktail. Lysates were mixed with 4× sample buffer and heated at 70 °C for 10 minutes. For VLPs samples, 1×10^10^ particles were directly mixed with 4× sample buffer and similarly heated at 70 °C for 10 minutes. All samples were resolved on NuPAGE™ 4-12% Bis-Tris Protein Gels (Thermo Scientific) at 120 V for 2 hours using NuPAGE™ MES SDS running buffer (Thermo Scientific). Proteins were transferred to membranes with the iBlot™ 2 Transfer Stacks (Thermo Scientific). Following transfer, membranes were blocked for 1 hour at room temperature with Intercept™ blocking buffer (LI-COR Biosciences) on a shaker, then incubated overnight at 4 °C with primary antibodies. The next day, membranes were washed three times with TBS-T (5 minutes each), followed by incubation with the appropriate secondary antibodies for 1 hour at room temperature under gentle agitation. After an additional three washes with TBS-T and one final wash with PBS, membranes were imaged using the Odyssey infrared imaging system (LI-COR). Uncropped blots are available either in the Source Data file (for main figures) or included at the end of the Supplementary Information (for Supplementary Figures). Comprehensive details on all primary and secondary antibodies used are listed in Supplementary Table 2.

### Flow cytometry

Conventional flow cytometry (MACSQuant 10, Miltenyi Biotec, Germany) was used to measure the GFP expression of the diverse traffic-light reporter cells/Stop-light reporter cells treated with Cre-loaded VLPs, Cas9 RNP (Cas12a RNP)-loaded VLPs at different time points, or GFP expression of the reporter cells co-cultured with VLPs-producing cells for 48 hours, or the GFP expression of B16F10-TL xenograft treated with Cre-loaded VLPs in vivo. Also, MACSQuant-based flow cytometry was used to measure the tdTomato expression of the primary cells from RS-26 mice treated with Cre-loaded VLPs ex vivo, or the primary cells derived from RS-26 reporter mice treated with engineered VLPs in vivo. Briefly, after the trypsinization, the cells in 96-well plates were resuspended in resuspended in 100 *μ*L of PBS containing 1.5% FBS, and 4ʹ,6-diamidino-2-phenylindole (DAPI). The cells were sampled by MACSQuant Analyzer 10 cytometer (Miltenyi Biotec, Germany). Data was analyzed with FlowJo software (version 10.8.1) and doublets were excluded by forward scatter area versus height gating (gating strategy available in Supplementary Fig. 7 and Supplementary Fig. 8). For spectral flow cytometry analysis of phenotypic cellular subsets in dissociated mouse organs, all samples, including unstained controls, single-stained controls and fluorescence-minus-one controls, were acquired on a Northern Lights spectral flow cytometer (Cytek) equipped with 405 nm, 488 nm, and 638 nm lasers. Mouse FcBlock reagent and Brilliant Stain Buffer Plus (both BD Biosciences) were added to samples according to manufacturer’s recommendations before the following antibodies were added: F4/80-BV510 (BD #743280, clone T45-2342), NK1.1-BV605 (Biolegend # 108753, clone PK136), CD4-BV650 (BD #563747, clone RM4-5), Ly6C-BV711 (BD #755195, clone HK1.4.rMAb), CD11b-BV750 (BD #746910, clone M1/70), CD45-BV786 (BD #564225, clone 30-F11), CD8a-PerCP-Cy5.5 (Biolegend #100734, clone 53-6.7), CD11c-PE/Cy7 (Biolegend #117318, clone N418), Ly6G-APC (BD #560599, clone 1A8), CD19-R718 (BD #562956, clone 1D3), and CD3-APCeFluor780 (eBioscience #47-0032-82, clone 17A2). DAPI was added for dead cell exclusion. Spectral unmixing was performed with SpectroFlo Software V3.3.0 (Cytek), and data was analysed, plotted and quantified with FlowJo software version 10.9.0 (FlowJo LLC). Gating strategy is provided in Supplemental Figure 9.

### Fluorescence microscopy

Following the treated with engineered VLPs or nanoblades, eGFP-positive cells and mCherry-positive cells were subsequently examined using fluorescence microscopy. Imaging fields were selected at random to minimize bias, and identical acquisition settings were applied across all experimental groups within a single experimental run. Post-acquisition image processing was performed uniformly, either directly through the imaging system software or using Fiji, to ensure consistency in data analysis.

### Mouse experiments

In the intra-tumor injection model, C57BL/6 mice were purchased at 5-week age and housed in our animal facility for at least one week before use according to standard routines (ambient temperature: 20–22 °C, humidity: 45–55%, dark/light cycle: 12/12 h). B16F10-TL cells were harvested and resuspended in PBS and then inoculate the cells subcutaneously at the number of 5×10^5^ per mouse. After 10 days of inoculation to form obvious tumors, engineered VLPs were injected directly into the tumors. The injected volume was 50 *µ*L per mouse with 2×10^11^ VLPs. Three days after intra-tumor injection of VLPs, the mice were sacrificed, and the tumors were harvested. Half of the tumor tissues were fixed with PFA to prepare the xenograft slides and the other half were dissociated into a single-cell suspension. The slides with tumoral tissue were stained to detect the GFP expression, and the single-cell suspensions were detected by the flow cytometry to measure the percentage of GFP positive cells in the whole xenograft. In the intracerebroventricular-minipump injection model, R26-LSL-tdTomato reporter mice were infused with particles (8×10^11^ VLPs per mouse) through intracerebroventricular (ICV) infusion to the CSF, as previous published^14^. Four days after infusion, the brains were harvested and detected by the flow cytometric analysis. In the systemic in vivo delivery study, R26-LSL-tdTomato reporter mice received intravenous injections of the particle formulations at a dose of 1×10^12^ particles per mouse. After 4 days, the mice were euthanized and sacrificed, and single-cell suspensions from liver, spleen, and bone marrow were prepared for flow cytometry analysis.

### Immunofluorescent staining for melanoma xenograft tissues

The slides were fixed for 1 hour at 65°C. After that, the slides were deparaffinized and rehydrated by immersing them in Xylene for 20 minutes, then 100% ethanol for 3 minutes twice, 95% ethanol for 3 minutes, 70% ethanol for 3 minutes, and 50% ethanol for 3 minutes. After that, the slides were washed with cold tap water for 5 minutes before antigen retrieval with citrate buffer, pH 6.0. (Sigma). Following antigen retrieval, the slides were washed three times in PBS for five minutes each. The tissues were then blocked for 30 minutes at 37°C using blocking buffer. After blocking, the slides were treated overnight with primary anti-GFP antibody (Abcam, ab290, 1:200 dilution). The primary antibody was preserved on the second day, after which it was washed three times in PBS for five minutes each. The slides were then incubated for 30 minutes at 37°C with the secondary antibody Goat Anti Rabbit IgG H&L (Alexa Fluor® 488) (Abcam, ab150077, 1:500 dilution), followed by three 5-minute washes with PBS. Finally, the slides were mounted with ProLongTM Diamond Antifade Mountant with DAPI (Thermo Scientific) and sealed with nail polish. Images were captured with a confocal fluorescent microscope (Nikon, Japan). Comprehensive details on all primary and secondary antibodies used are listed in Supplementary Table 2.

### H&E staining of liver

Sagittal brain sections (5 µm) were processed for routine H&E histology. Slides were deparaffinized in xylene, rehydrated through graded ethanols to water, stained with hematoxylin, rinsed in tap water, counterstained with eosin, then dehydrated and cleared in ethanol/xylene before coverslipping. Representative regions of liver were imaged by light microscopy and reviewed by an experienced pathologist.

## Statistics

Results are provided as mean (± standard deviation) of biological replicates where applicable. Two-sided Student’s t-test was used to compare the difference between two groups. multiple groups with a single independent variable were analysed using one-way ANOVA or using two-way ANOVA for multiple groups with multiple independent variables. All collected data were analyzed with no exclusions. Mice and samples were randomly allocated to experimental groups. The Investigators were not blinded to allocation during experiments and outcome assessment. * *p* < 0.05, ** *p* < 0.01, *** *p* < 0.001 and **** *p* < 0.0001 were set as statistical significance levels.

## Data availability

All the data generated in this study are provided in the Supplementary Information. Materials are available through material transfer agreement (MTA) submitted to S.E.-A. and Evox Therapeutics Limited, Oxford, United Kingdom.

## Acknowledgements

This study was supported by the Swedish Research Council (VR, grant 2021-02407) for a half-time clinical research position and a CIMED junior investigator grant awarded to J.Z.N.; by the Swedish Research Council (VR, grant 2022-02449) for a half-time clinical research position and a CIMED junior investigator grant (FoUI-976434) awarded to O.W.; by a CIMED junior investigator grant (FoUI-988637) awarded to Xiuming Liang; and by Cancerfonden (211762 Pj 01 H), the European Research Council (ERC) under the EU Horizon 2020 research and innovation programme (DELIVER, grant No. 101001374; EXPERT, grant No. 825828), the Swedish Foundation for Strategic Research (FormulaEx, SM19-0007), the Brain Foundation (Hjärnfonden, contract FO2024-0073-TK-113), and the Swedish Research Council (4–258/2021) awarded to S.E.-A. Additional support was provided by Evox Therapeutics Limited.

Histological analysis as well as FFPE sample preparation was performed at the P Morphological Phenotype Analysis (FENO), at Karolinska Institutet (Sweden), supported by Karolinska Forskning och Utbildning, Utveckling (FoUU) and Karolinska Institutet infrastructure council.

## Author contributions

Conceptualization: G.Z., S.E.-A., and X.L. Methodology: G.Z., V.W.Q.H., H.Z., S.R., J.S., O.G., Z.N., D.R.M., L.V.H., O.W., and A.G. Investigation: G.Z., X.L., V.W.Q.H., H.Z., S.R., J.S., O.G., Z.N., D.R.M., R.E.V., O.W., J.Z.N., A.G., and S.E.-A. Visualization: G.Z., X.L., and S.E.-A. Funding acquisition: J.Z.N., O.W., X.L., and S.E.-A. Supervision: X.L., and S.E.-A. Writing: G.Z., and X.L. Editing: X.L., A.G., and S.E.-A.

## Competing interests

O.W., J.Z.N., A.G., and S.E.-A. serve as consultants and stakeholders in Evox Therapeutics Limited, Oxford, United Kingdom. All data supporting this study are provided in the manuscript and Supplementary Information. The other authors declare no competing interests.

