## Supplementary figures for "Efficient delivery of gene editors using intein-engineered virus-like particles"

**Supplementary materials**

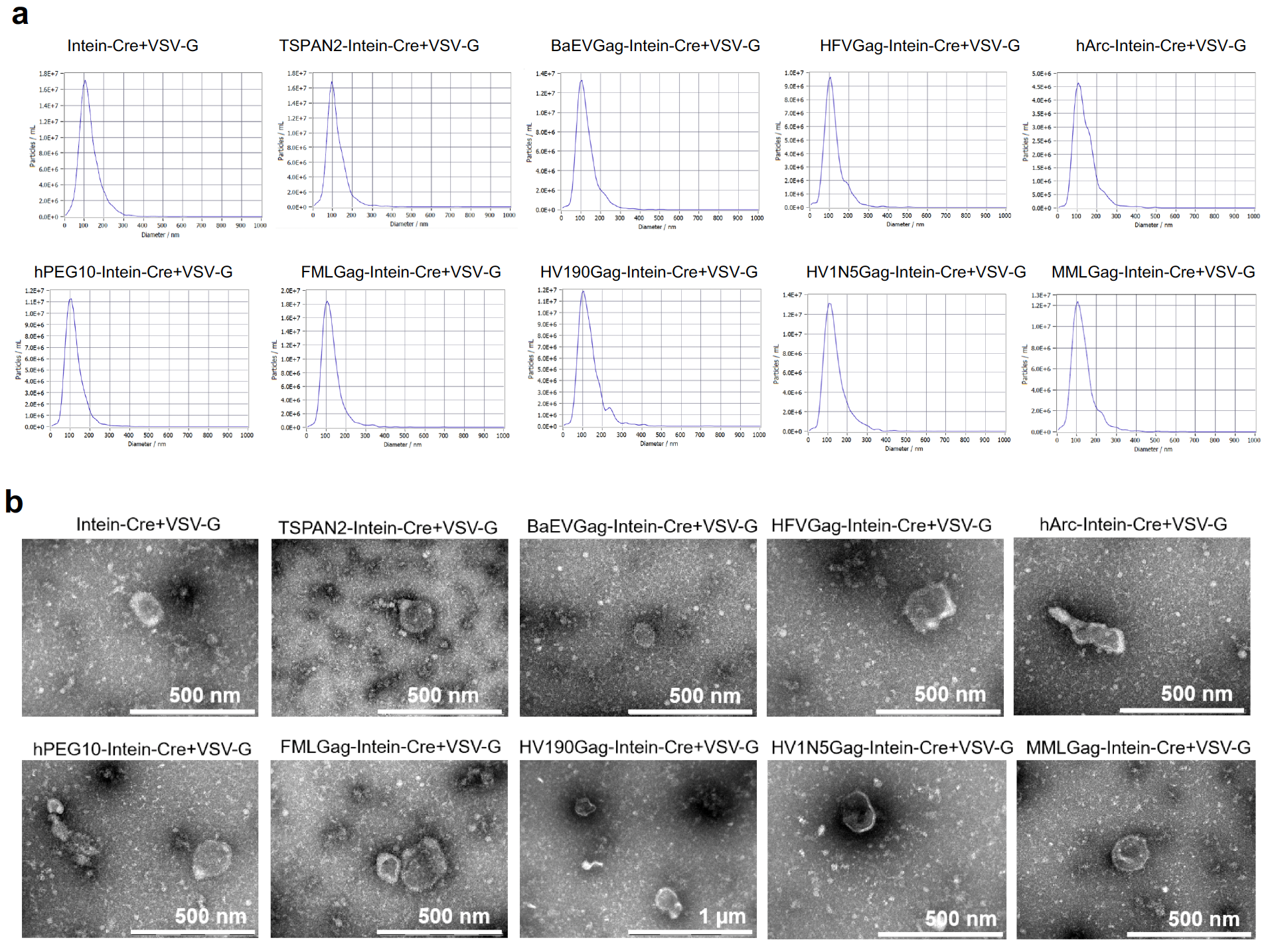

**Supplementary Fig 1. Characterization of intein-engineered particles generated from different Cre-delivering constructs.** **(a)** Transmission electron microscopy (TEM) images of negatively stained particles produced by transfecting HEK293T cells with various Intein–Cre fusion constructs co-expressed with VSV-G. All constructs include different sorting scaffolds such as TSPAN2, BaEV-Gag, HFV-Gag, hArc, hPEG10, FMLV-Gag, HV190Gag, HIV1NS-Gag, MMLV-Gag, or Intein alone. scale bar: as indicated. **(b)** Size distribution profiles of the corresponding particle preparations measured by nanoparticle tracking analysis (NTA) using a ZetaView instrument. All samples show characteristic EV-like size ranges (average size peak at around 90 nm to120 nm).

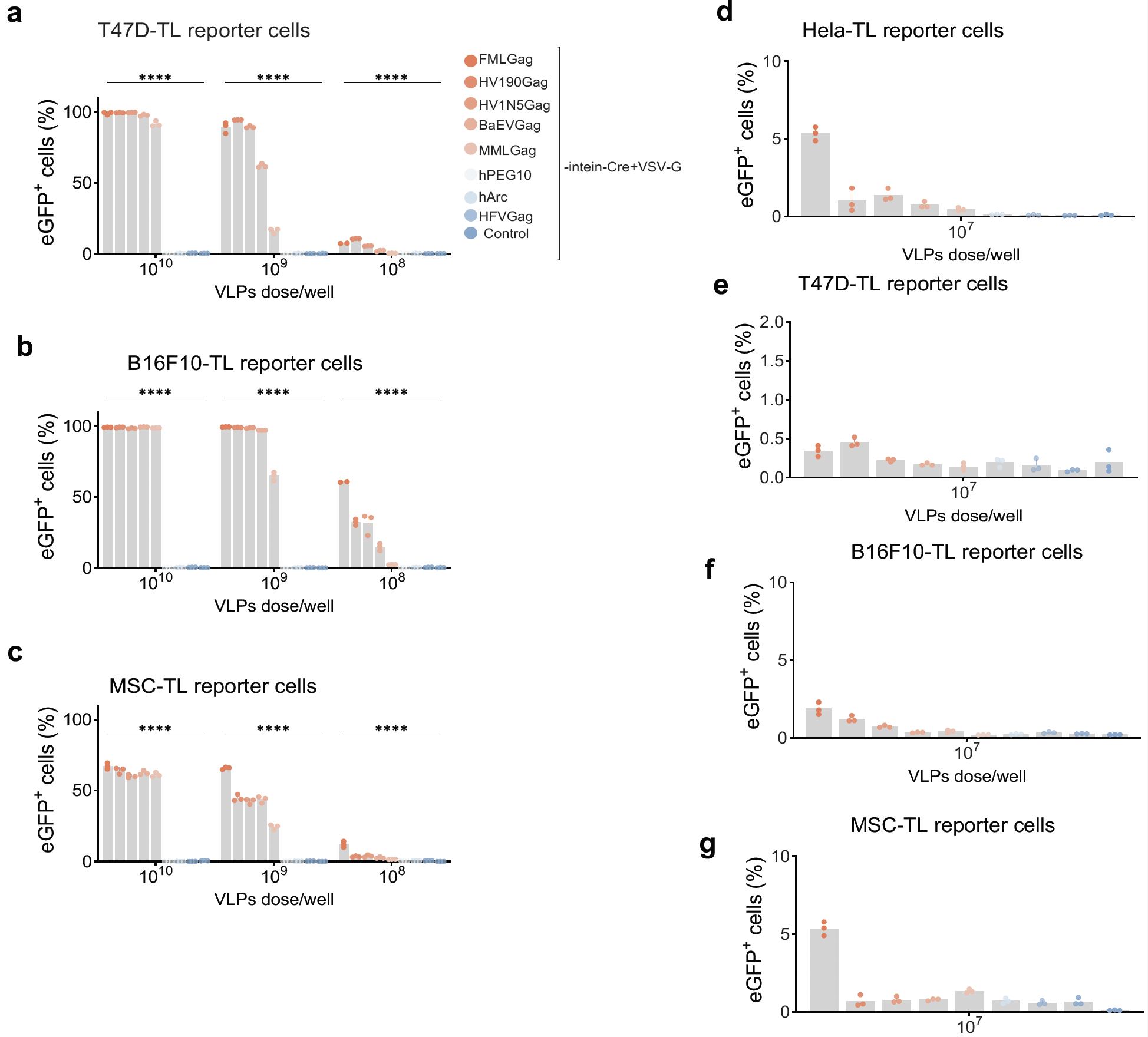

**Supplementary Fig. 2 Identification of high-efficiency intein-engineered VLPs scaffold proteins for functional intraluminal cargo delivery across a range of VLP doses.** **(a-c)** Quantification of eGFP⁺ cells in T47D-TL (a), B16F10-TL (b), and MSC-TL (c) reporter cell lines after 48 h incubation with Cre-loaded VLPs engineered using various Gag or EV-related scaffold proteins with indicated doses. Control group is the Cre-loaded particles without VSV-G pseudo typing. **(d–g)** Quantification of eGFP⁺ cells in 4 TL reporter cell lines (HeLa-TL, T47D-TL, B16F10-TL, and MSC-TL) after 48 h incubation with Cre-loaded VLPs engineered using various Gag or EV-related scaffold proteins. Bar graphs on the left show dose-dependent functional delivery; right panels specifically highlight delivery efficiency at the ultra-low dose of 1×10^7^ particles/well. Data are presented as mean ± SD. *****p < 0.0001*.

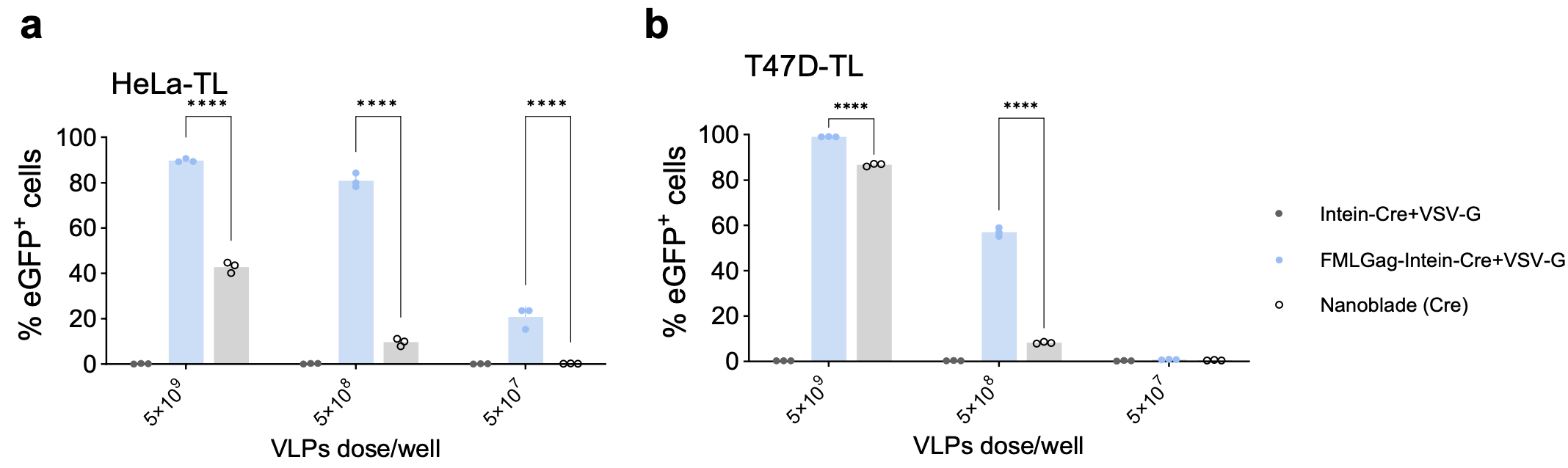

**Supplementary Fig 3. Comparison of intein-engineered VLP-based Cre delivery with Nanoblade system at different timepoints and doses across cell lines.** **(a–b)** Percentage of eGFP⁺ cells in HeLa-TL(a) and T47D-TL (b) reporter cells respectively after treatment with different Cre-delivering particle formulations at three indicated doses. Flow cytometry analysis at 48 h post-treatment. Data are presented as mean ± SD. *****p < 0.0001*.

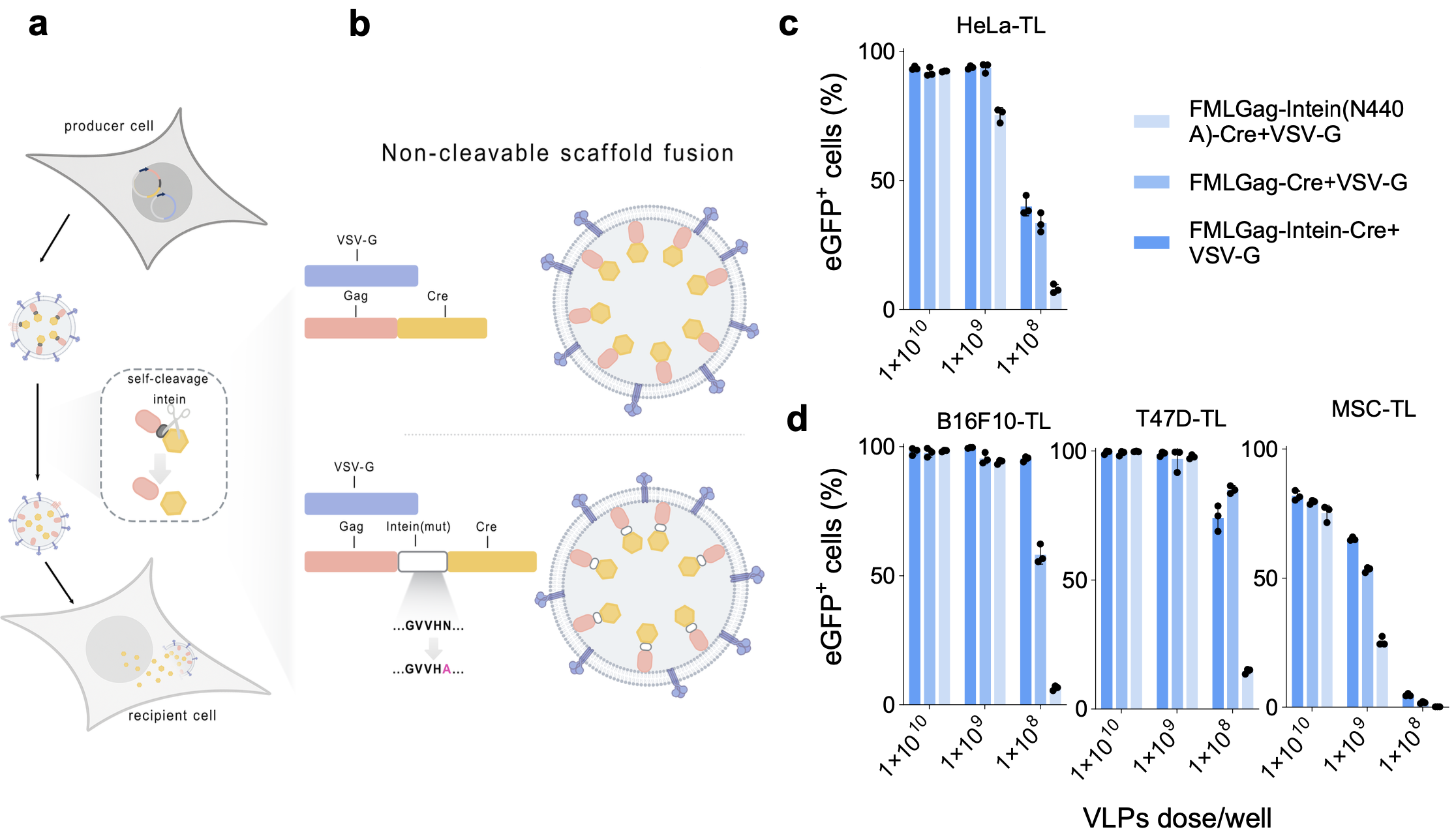

**Supplementary Fig 4. Engineering strategies for VLPs-mediated cargo delivery.** (**a**) Schematic overview of engineering VLPs using intein. (**b**) Non-cleavable scaffold fusion, in which the intein domain is either deleted or inactivated (N440A mutation), resulting in direct fusion of Cre to the scaffold. (**c-d**) Flow cytometry analysis of eGFP⁺ cells in TL reporter lines (HeLa-TL, B16F10-TL, T47D-TL, and MSC-TL) 48 h after exposure to VLPs generated with different engineering strategies at serial doses. Data are presented as mean ± SD

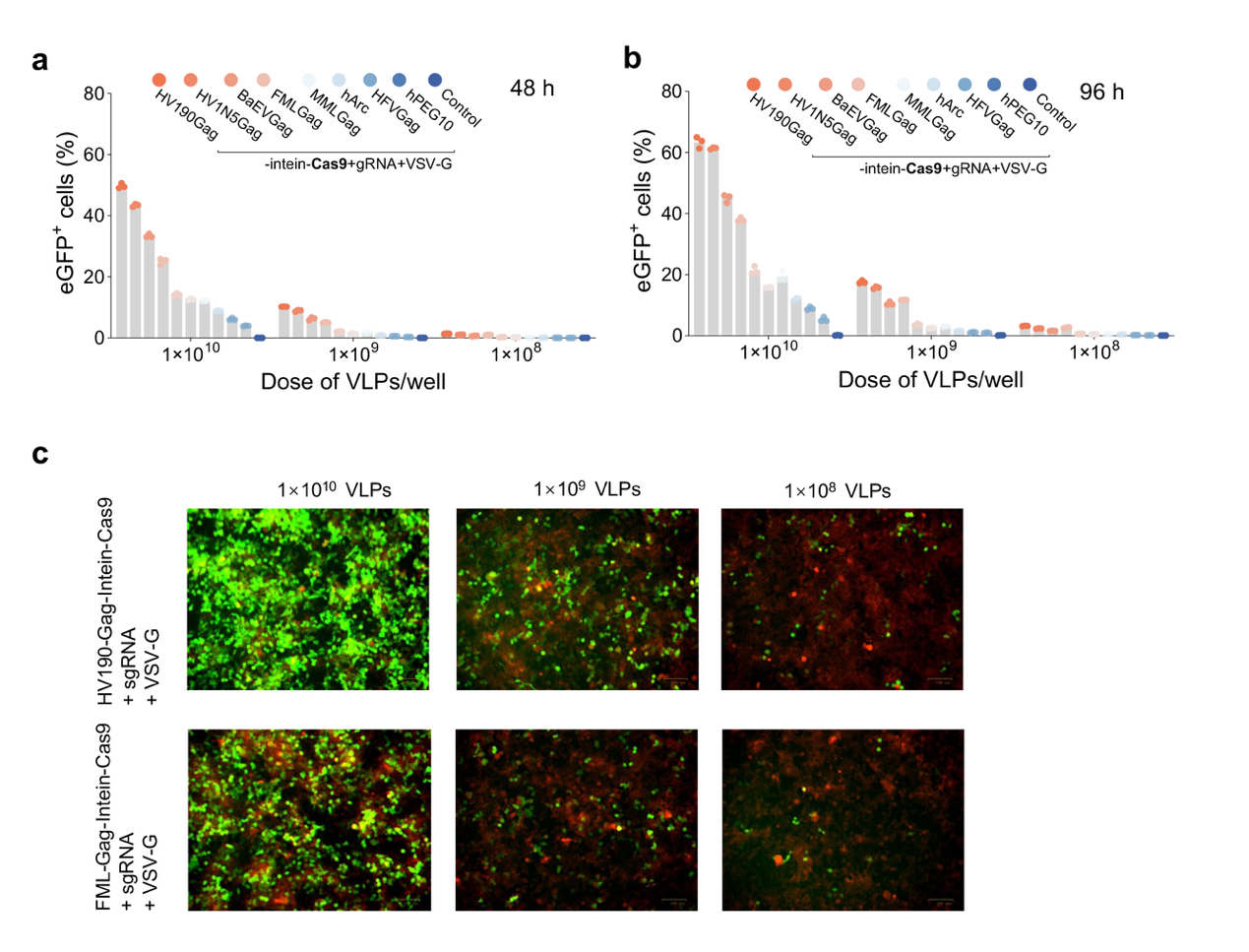

**Supplementary Fig. 5 Time- and dose-dependent analysis of Cas9 RNP delivery via intein-engineered VLPs.** **(a–b)** Percentage of eGFP⁺ cells measured by flow cytometry at 48 h (a) and 96 h (b) post-treatment. Control group is the Cas9 RNP-loaded particles without VSV-G pseudo typing. **(c)** Representative immunofluorescence images of HEK-SL cells treated with Cas9 RNP VLPs at varying doses. eGFP expression indicated successful gene editing. Scale bar, 100 μm. Data are shown as mean ± SD.

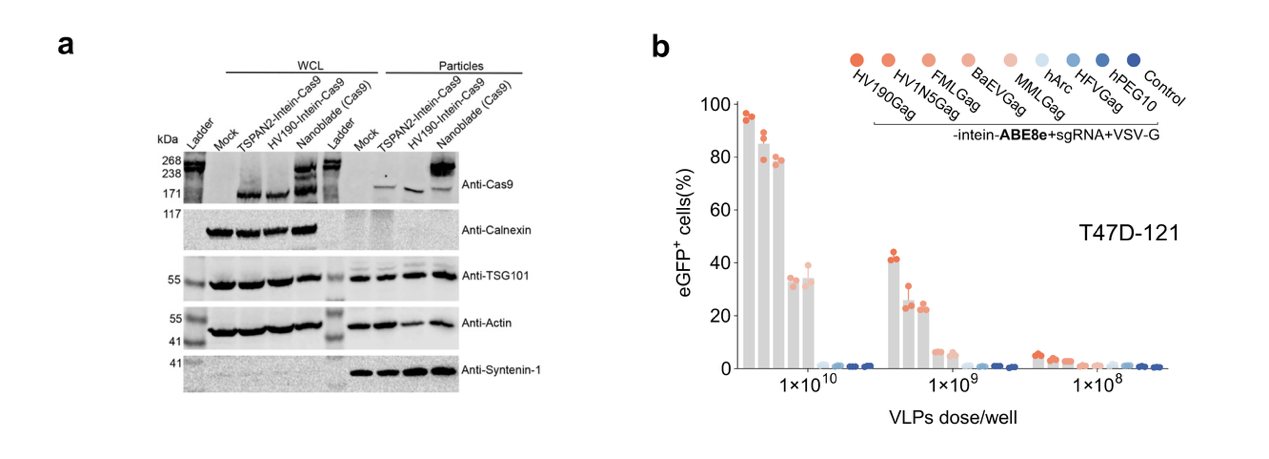
**Supplementary Fig. 6 Western blot validation of intein-engineered VLPs packaging and quantification of base editing efficiency. (a)** Western blot was conducted with lysates from 5×10^5^ particle-producing cells and 1×10^10^ engineered particles respectively. TSG101, syntenin-1, and β-actin were used as particle markers, while calnexin (an endoplasmic reticulum marker) was included to verify the absence of cellular contamination in particle samples. (**b**) Percentage of eGFP+ cells in T47D-121 reporter cells 72 h after post-treatment with Base Editor (ABE8e) RNP VLPs produced using different sorting domains. Control group is the base editor-loaded particles without VSV-G pseudo typing. Data are presented as mean ± SD.

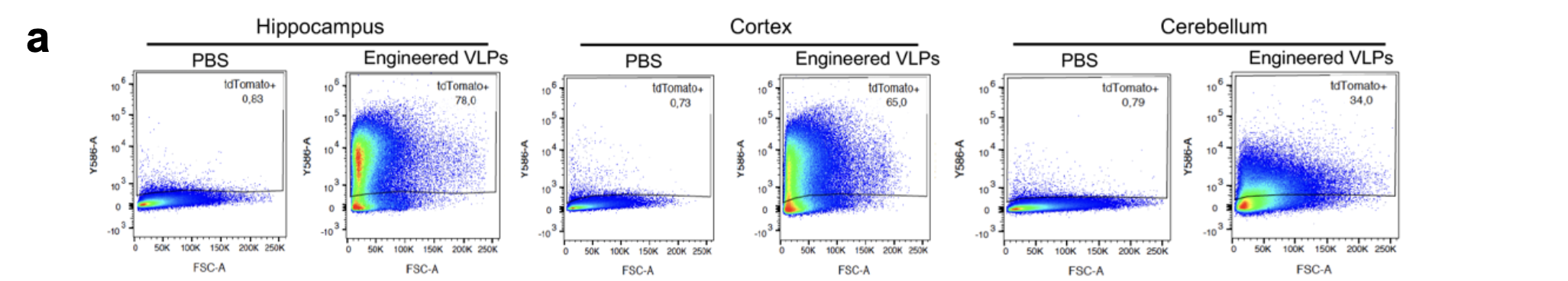

**Supplementary Fig. 7 Representative images showing flow cytometry analysis of tdTomato expression in hippocampus, cortex, and cerebellum after ICV minipump delivery of Cre-loaded particles.** n = 3 mice for engineered particle group and n = 2 mice for PBS group.

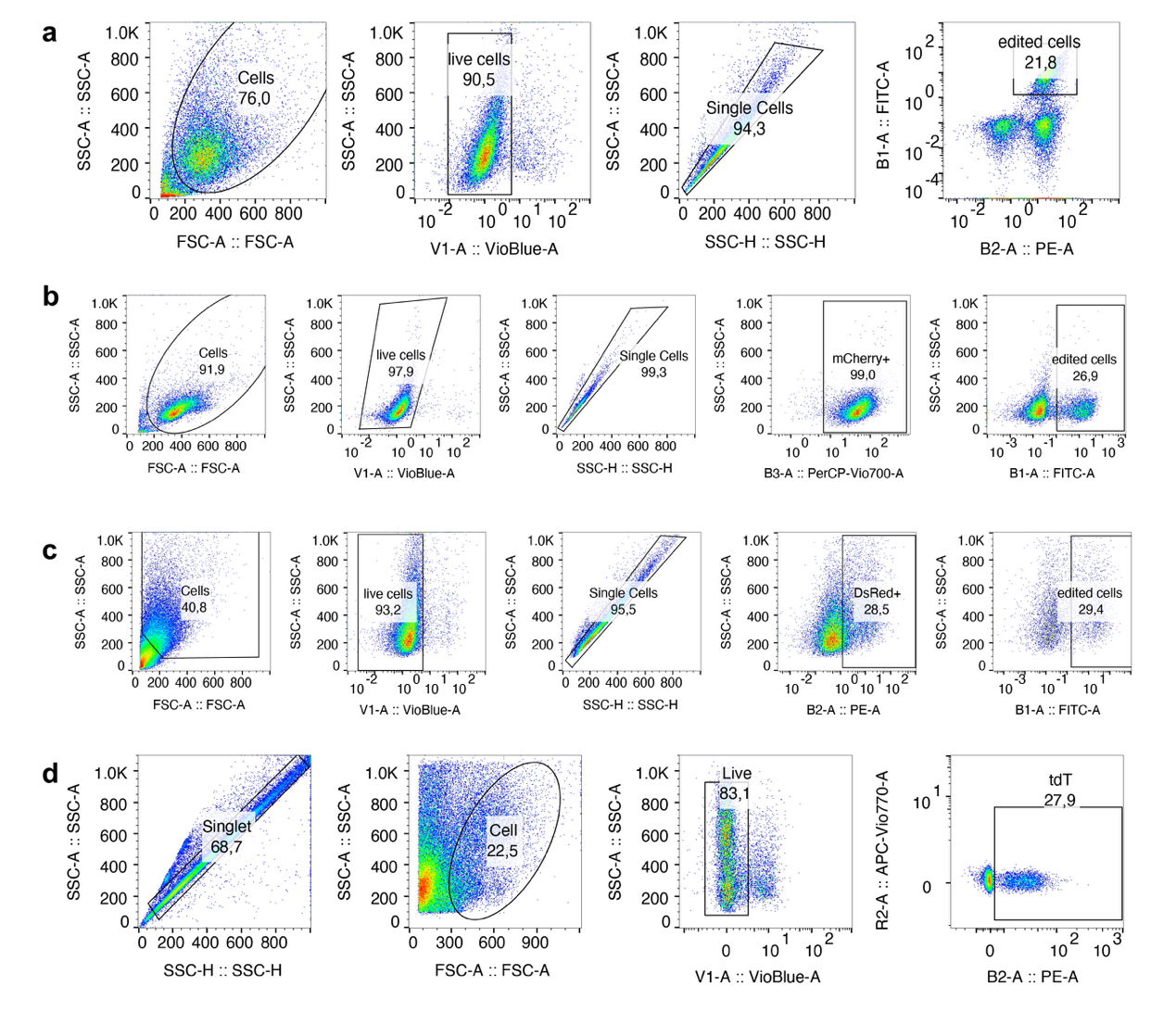

**Supplementary Fig. 8** **Gating strategy for flow cytometric analysis in this study.** (**a**) Related to Figure 1e-1f, Figure 2d-2g, Supplementary Figure 2, Supplementary Figure 3 and Supplementary Figure 4. (**b**) Related to Figure 2d, 2e, Supplementary Figure 5a-5b. (**c**) Related to Figure 4c. (**d**) Related to Figure 3c-3e, and 3f.

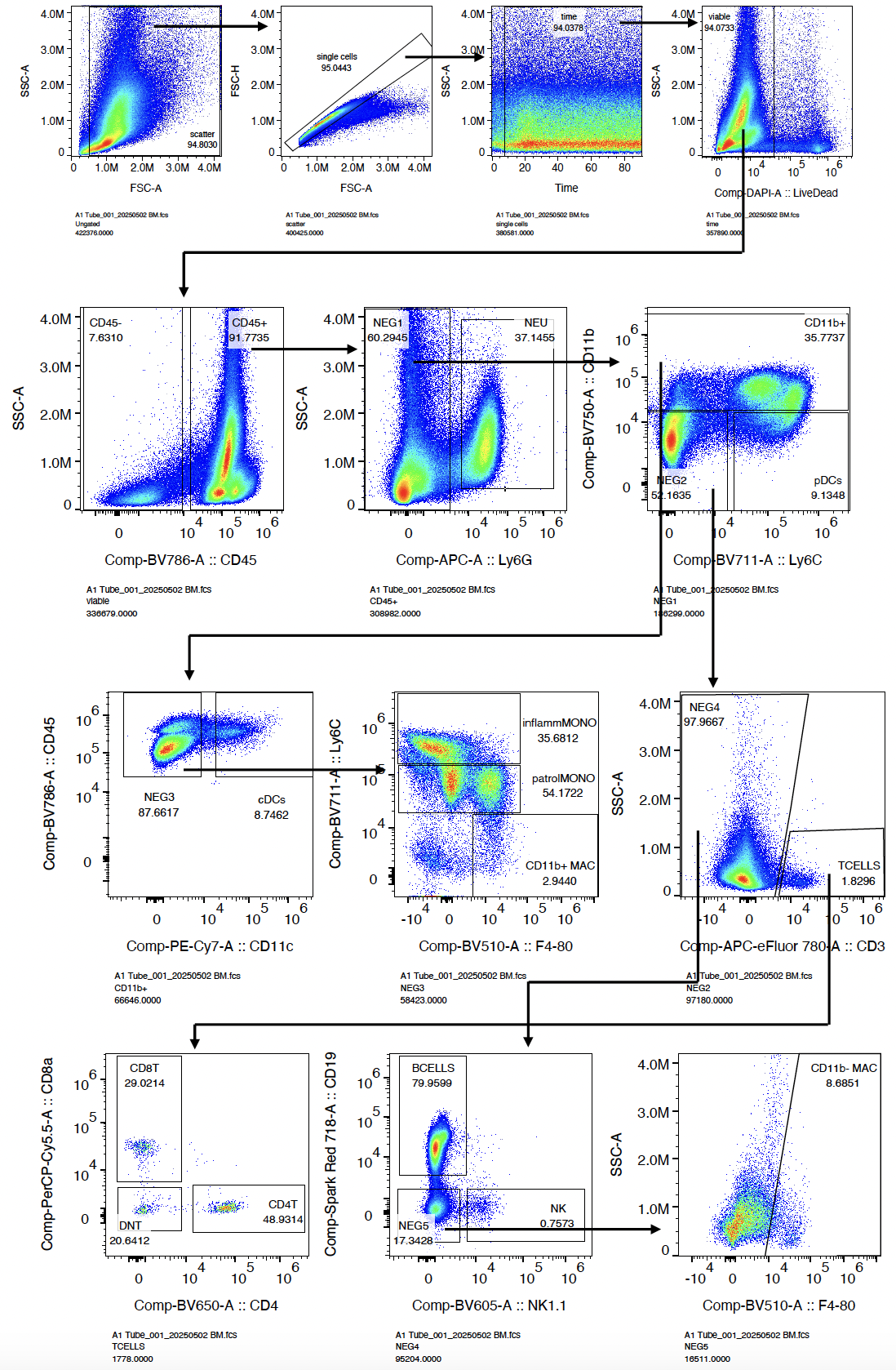

**Supplementary Fig. 9** **Gating strategy for identification of cellular subsets in this study by spectral flow cytometry analysis.** Related to Figure 5b-5d. Data shows example gating for a bone marrow sample for identification of hematopoietic (CD45^+^) versus non-hematpoietic (CD45^-^) cells as well as Neutrophils (NEU), plasmacytoid dendritic cells (pDCs), conventional dendritic cells (cDCs), inflammatory (inflammMONO) and patrolling (patrolMONO) monocytes, CD11b^+^ macrophages (CD11b^+^ MAC), T cells (TCELLS), CD4 (CD4T) and CD8 (CD8T) T cells, natural killer cells (NK), B cells (BCELLS), and CD11b^-^ macrophages (CD11b^-^ MAC).

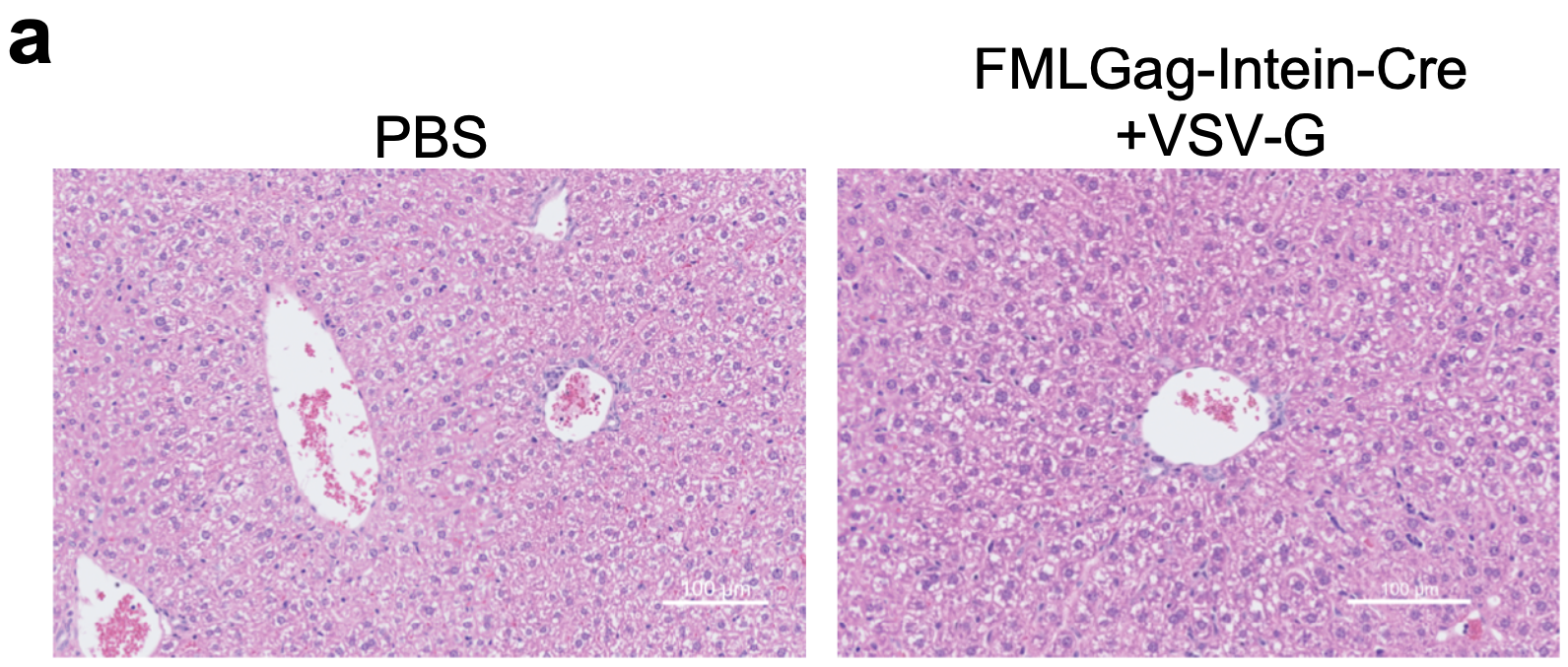

**Supplementary Fig. 10 Representative histology images of liver after 4 days of particle injection intravenously.** Scale bar, 100 μm.
